# Are reading and face processing related? An investigation of reading in developmental prosopagnosia

**DOI:** 10.1101/039065

**Authors:** Randi Starrfelt, Solja K. Klargaard, Anders Petersen, Christian Gerlach

**Author notes:** Corresponding author: Randi Starrfelt Dept. of Psychology University of Copenhagen O Farimagsgade 2a DK-1351 Copenhagen K Denmark.

## Abstract

Recent hypotheses suggest that learning to read influences the cognitive and cerebral organization of other perceptual skills, including face processing. Developmental prosopagnosia (DP) is a disorder of face recognition in the absence of acquired brain injury, and in the context of normal intelligence and general cognitive development. To shed light on the potential relationship between reading and face processing in this group (and in general), we investigated reading performance in 10 participants with DP and 20 matched controls. We find that the group of DPs perform strikingly similar to the control group on four sensitive reading tests measuring visual recognition and naming of single letters and words, word length effects, and text reading speed and comprehension. Thus, there is a clear dissociation between impaired face processing and preserved reading in this group, a finding that challenges the recently proposed hypothesis that reading acquisition and face processing abilities are intrinsically linked. Developmental prosopagnosics can learn to read as fluently as normal subjects, while they are seemingly unable to learn efficient strategies for recognizing faces.

## Introduction

*Developmental dyslexia* is a well-known syndrome characterized by impaired ability to learn to read and spell. The syndrome *Developmental prosopagnosia (DP)*, on the other hand, has received comparatively less attention. DP is a hereditary disorder, estimated to affect 2 - 2.5% of the population (Bowles et al., 2009; Kennerknecht et al., 2006). People suffering from DP fail to develop normal face recognition abilities, and can have difficulties even in recognizing their own immediate family. Many readers will be familiar with Oliver Sacks’ report of “The man who mistook his wife for a hat” (Sacks, 1985), describing a man suffering from visual agnosia and prosopagnosia caused by a brain disease. Fewer will know that Oliver Sacks himself suffered from a deficit in face recognition, one that was present throughout his life, and with no known neurological cause: developmental prosopagnosia (Sacks, 2010).

Similar to these developmental disorders, acquired deficits in reading and face recognition also exist. Deficits in reading and face recognition can result from injury to posterior brain areas, and in some cases, these deficits appear quite selective. In *pure alexia*, visual word processing is impaired, while identification of faces may be relatively spared, and in acquired *prosopagnosia*, face recognition is impaired, while reading is commonly reported to be unaffected. This constitutes a double dissociation, indicating that words and faces are processed by different mechanisms and brain areas; or so textbook knowledge holds (Gazzaniga, Ivry, & Mangun, 2013). Processing of words and faces has been linked to specialized perceptual brain areas lateralized to different hemispheres; the visual word form area (VWFA; Cohen et al., 2000) in the left fusiform gyrus, and the fusiform face area (FFA; Kanwisher, McDermott, & Chun, 1997) in the right hemisphere. This double dissociation between reading and face processing, and the modular theories of perception that often go with it, has recently been challenged by studies suggesting a relationship between reading and face recognition (Behrmann & Plaut, 2013; Cantlon, Pinel, Dehaene, & Pelphrey, 2011; Dehaene et al., 2010; Dundas, Plaut, & Behrmann, 2013, 2014). Perhaps the most intriguing results from this line of evidence come from studies showing that learning to read affects the cerebral substrate for face processing in a systematic way, and may even drive the lateralization of face processing to the right hemisphere (Dehaene & Cohen, 2007, 2011; Dundas et al., 2013).

If learning to read affects the lateralization of, and maybe even the cerebral specialization for face processing, what happens if face processing has not developed normally when children learn to read? Does such a developmental deficit in face processing affect the ability to learn to read, and if so, in what way? This has, to our knowledge, never been systematically examined.

To investigate this, we have tested 10 subjects with developmental prosopagnosia and matched controls on tests of letter, word, and text reading. If face processing and reading are dissociable processes, we would expect the group of developmental prosopagnosics to perform normally on reading tests. If, on the other hand, a normal development of the face recognition network is necessary for normal reading acquisition, then reading may be affected in developmental prosopagnosia. To ensure that reading skills were measured sensitively, we apply both experimental and psychophysical tests of letter and word recognition, as well as a more standard test of text reading.

## General Method

### Participants

All participants provided written informed consent according to the Helsinki declaration to participate in the project. The Regional Committee for Health Research Ethics in Southern Denmark has assessed the project, and found that it did not need formal registration.

#### Participants with developmental prosopagnosia (DPs)

Following appearances in Danish media, where we have informed about developmental prosopagnosia, we have been contacted by a number of people complaining of face recognition problems. They all report difficulties recognizing friends, colleagues, and sometimes even close family members and themselves by their faces, and that these problems have been present throughout their life.

10 of these subjects are included in the current study. As there are no established diagnostic criteria for DP we initially included subjects in our sample if they performed abnormally on the Cambridge Face Memory Test (CFMT; Duchaine & Nakayama, 2006) and/or the Cambridge Face Perception Test (CFPT; Duchaine, Germine, & Nakayama, 2007) when compared to the age and gender adjusted norms provided by Bowles et al. (2009). These tests were kindly provided by Brad Duchaine and translated into Danish. In the current study, only participants who performed more than 2SD’s below the mean of the matched control group on the CFMT, the most commonly used diagnostic test, were included. All included DP’s also report severe difficulties with face recognition in their everyday life, as evaluated by the face recognition part (29 items) of the Faces and Emotion Questionnaire (FEQ; Freeman, Palermo, & Brock, 2015)The questionnaire was kindly provided by Romina Palermo and Jon Brock, and translated to Danish by the last author. Three DP subjects were left handed, and all performed within the normal range on The Autism-Spectrum Quotient (AQ) questionnaire (Baron-Cohen, Wheelwright, Skinner, Martin, & Clubley, 2001). See Table 1 for an overview of age, gender, and basic test scores. The DP subjects did not receive remuneration for their participation in this study. The DPs (and controls) included in this study have participated in extensive testing in our lab, and we report here all tests where reading is measured.

For the DPs to be anonymous, and yet recognizable across publications, we have kept their project subject-numbers in the text and tables.

**Table 1.** Age, gender and performance (raw scores) on the Cambridge Face Memory Test (CFMT), the Cambridge Face Perception Test (CFPT), and the Face recognition questionnaire (FEQ) for the 10 participants with developmental prosopagnosia, and the mean and SD for the controls’ scores on these tests. Values in boldface designate performance deviating more than 2 SDs from the mean of the matched control group. In the CFMT a low score indicates a deficit, while in the CFPT and in the FEQ a high score indicates a deficit. The maximum score on the FEQ is 87.

| Subject | Age | Gender | Handedness | CFMT | CFPT | FEQ |
| --- | --- | --- | --- | --- | --- | --- |
| PP04 | 57 | M | Right | **37** | **86** | **71** |
| PP07 | 40 | F | Right | **41** | 60 | **66** |
| PP09 | 40 | F | Left | **43** | **70** | **52** |
| PP10 | 34 | F | Right | **33** | 58 | **62** |
| PP13 | 51 | M | Right | **35** | 42 | **64** |
| PP16 | 23 | F | Left | **39** | **64** | **54** |
| PP17 | 49 | F | Right | **35** | **88** | **56** |
| PP18 | 38 | F | Left | **30** | **78** | **69** |
| PP19 | 16 | M | Right | **33** | 48 | **53** |
| PP27 | 25 | M | Right | **42** | **66** | **59** |
| Control mean (SD) |  |  |  | 59.1 (7.9) | 41.3 (11.4) | 22.4 (11.4) |

#### Control subjects

We compared the 10 DPs with 20 control subjects; two matched on age, gender, and educational level to each individual with DP (three left handed). Thus the groups were comparable in terms of age (DP *M* = 37, range = 16-57; Control *M* = 37, range = 16-56) and years of education (DP *M*= 15.5, range = 11-17; Control *M* = 15.2, range = 10-17). All controls performed within the normal range on the CFPT and the CFMT, evaluated by the Bowles (2009) norms. Controls received gift certificates of ∼120 DKK (∼20 USD) per hour for their participation.

The statistical analyses below are based on principles of “New statistics” (Cumming, 2014). Individual data for the DPs in all experiments are presented in Supplemental table S1.

## Experimental Investigation

### Experiment 1. Reading latency and word length effects

While the reading latency of normal readers is little affected by the number of letters in a word, a *word length effect* is a common symptom of reading disorders (Barton, Hanif, Björnström, & Hills, 2014), both developmental and acquired. In pure alexia, RTs may increase with hundreds of milliseconds per additional letter in a word. Word length effects are also characteristic in beginning readers, and a drastic reduction of this effect characterizes successful reading acquisition.

#### Stimuli and procedure

In this experiment, we tested whether our sample with DP showed a word length effect (WLE) in single word reading. The exact paradigm has previously revealed elevated RTs and WLEs in patients with pure alexia (Habekost, Petersen, Behrmann, & Starrfelt, 2014; Starrfelt, Nielsen, Habekost, & Andersen, 2013). Stimuli were 150 words of 5–7 letters (50 of each length matched for word frequency and neighbourhood-size). Reading RTs were measured by a voice key (a microphone connected to the response box). The WLE is calculated using linear regression, where the slope represents the additional time needed per additional letter in a word. Mean overall RT was also calculated for each subject.

#### Results

Voice key errors (setting of the microphone too early /late) were excluded from the analysis. RTs were analysed for correct trials only. RTs were trimmed by excluding RTs from trials deviating more than 2.5 SD for each individual at each word length. On average 4.9% (range: 1.3 -7.3) of the trials for each DP was removed due to voice key errors or trimming. For the control participants it was 3.3% (range: 0.7 - 6).

The DP-group made on average 1% errors (range: 0 - 5%) whereas the control group made on average 0.5% errors (range: 0 - 3%). The mean correct reading RT for the DP group was 569 ms (95% CI = [519, 619]) and 521ms (95% CI = [489, 553]) for controls. The mean WLE for the DP group was 10 ms (95% CI = [2, 19]) and 11 ms (95% CI = [6, 16]) for controls. As can be seen from Figure 1 there are overlaps in the CIs of the mean reading RTs for the DPs and controls, and the main parameter of interest in this test, the WLE, is strikingly similar for the two groups.

**Figure 1.**
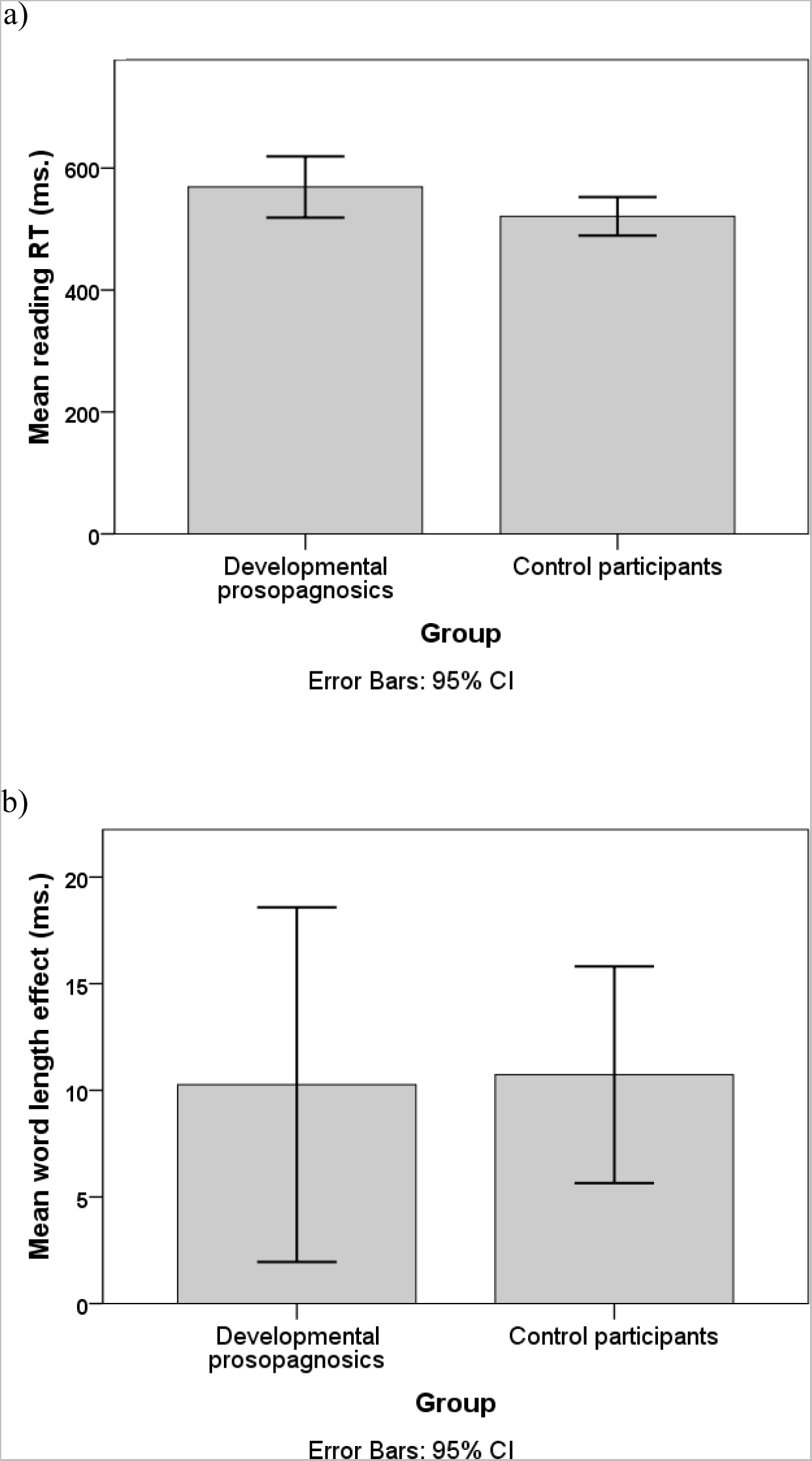
Mean overall word reading time (panel a) and mean word length effect (panel b) for developmental prosopagnosics (*N* = 10) and control participants (*N* = 20).

### Experiment 2. Word superiority effect

Fluent reading is characterised by fast and parallel processing of letters in words; a likely explanation for the minimal WLE in proficient readers. In fact, normal readers typically identify letters in words faster than they identify single letters, a phenomenon known as the *word superiority effect* (WSE) (Reicher, 1969; Wheeler, 1970). We have recently developed a psychophysical paradigm for measuring the WSE comparing short words and single letters, and found that the effect is robust over a range of exposure durations in healthy participants (Starrfelt, Petersen, & Vangkilde, 2013). In addition, and more surprisingly, we also found a WSE in vocal reaction times to stimuli presented in free viewing conditions; words were named faster than singly presented letters. Patients with pure alexia, which is thought to be caused by a breakdown in parallel letter processing, do not show a WSE in either of these tasks (Habekost et al., 2014).

#### Stimuli and procedure

This paradigm consists of two experiments, an RT task (Experiment 2a) and a psychophysical task using limited exposure durations (Experiment 2b), both described in detail in Starrfelt, Petersen, et al. (2013; Experiments 1 and 2). The stimuli are the same in both the RT and the psychophysical task; 25 letters (w excluded), and 25 three letter words. The words are confusable in the sense that none can be identified uniquely by seeing one letter only, and for most of the words all three letters must be processed for the word to be correctly identified (see Appendix in Starrfelt, Petersen, et al., 2013).

#### Experiment 2a. Letter and word naming

This is a computerised naming task, and the procedure is similar to Experiment 1. The purpose of this RT-task was to familiarise subjects with the word stimuli employed in the psychophysical paradigm (Experiment 2b). Everyone knows the letters of the alphabet, and we wanted to ensure that the subjects also knew which words were used as stimuli. The letter and word conditions included 50 trials each, and were run separately, the letter task first. The task was to name the word or letter presented on the screen. There were ten practice trials in each condition.

#### Results

Voice key and naming errors were recorded, and excluded from the RT analysis. RTs were trimmed for each participant by excluding RTs from trials deviating more than 2.5 SD from the individual mean. 2.9% (range: 0 - 8) and 1.3% (range: 0 - 4) of the trials for each DP was removed due to voice key errors or trimming for letters and words respectively. For the control participants it was 1.2% (range: 0 - 4) and 0.6% (range: 0 - 4) respectively. 9 DPs participated (PP16 did not and accordingly, the two controls matched to PP16 were also excluded from the analyses).

The DP-group made no errors. The controls made on average 0.9% errors with letters (range: 0 - 6%) and 0.4% errors with words (range: 0 - 4%). For the DPs the mean RT to letters was 488 ms (95% CI = [450, 526]) and 451 ms (95% CI = [417, 484]) to words. For controls the mean RT to letters was 473 ms (95% CI = [452, 495]) and 466 ms (95% CI = [446, 486]) to words. As can be seen in Figure 2 there is again considerable overlap in the CIs for the DPs and the controls for both letter and word naming, suggesting that they perform quite similarly in this experiment. Note, however, that the difference between letter and word naming RTs (the WSE) is rather modest for the controls (*d_z_* = 0.21; 95% CI *Mdif* = [-11, 25]) in comparison with that of the DPs (*d_z_* = 1.02; 95% CI *M_dif_* = [9, 65]). This difference is seen more clearly in Figure 3, which depicts the WSE in RT (the mean difference in RT to words compared to letters) where the mean of this difference for the DPs lies considerably outside the 95% CI of the mean difference for the controls.

**Figure 2.**
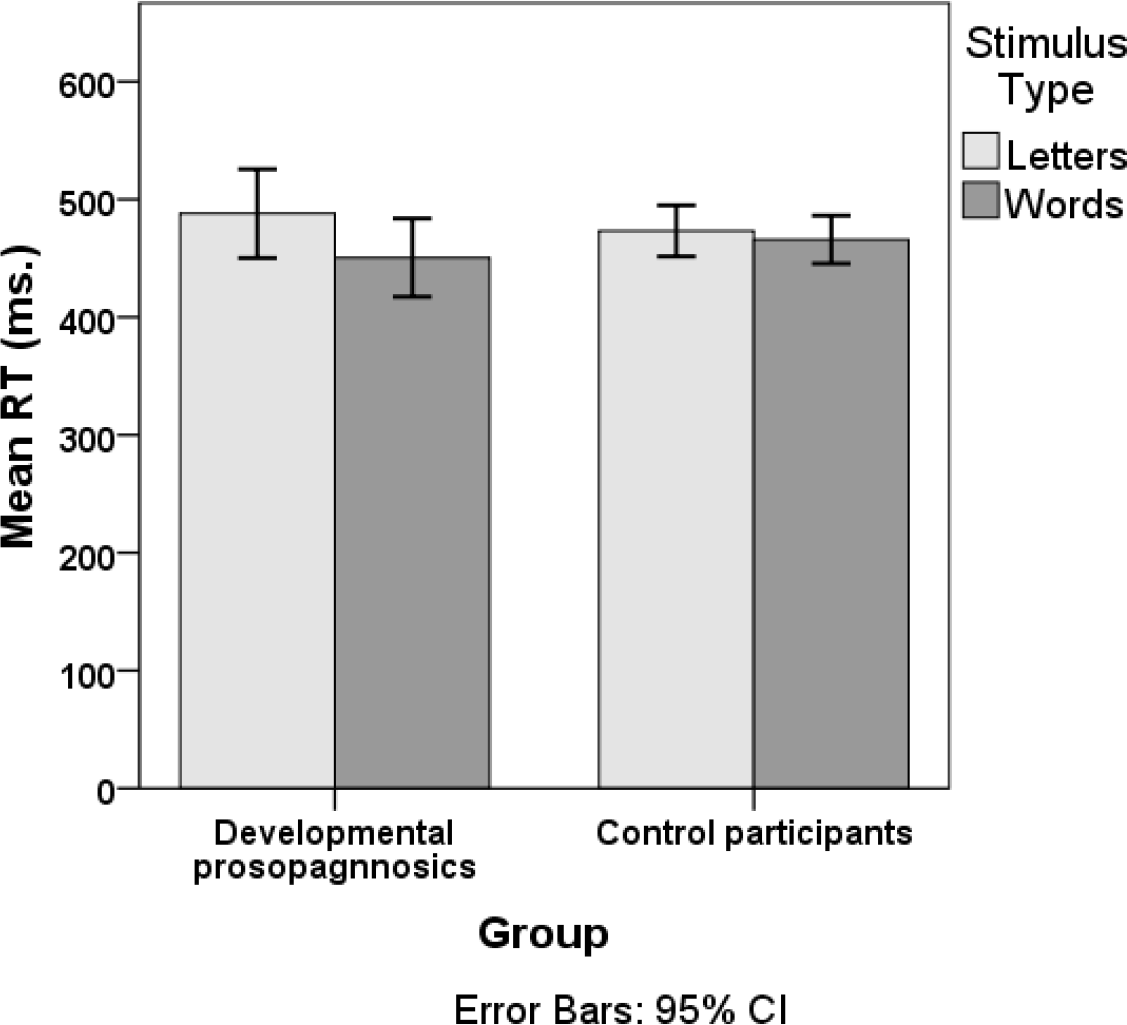
Naming time. Mean RTs to single letters and words in Experiment 2a for DPs (*n* = 9) and controls (*n* =18).

**Figure 3.**
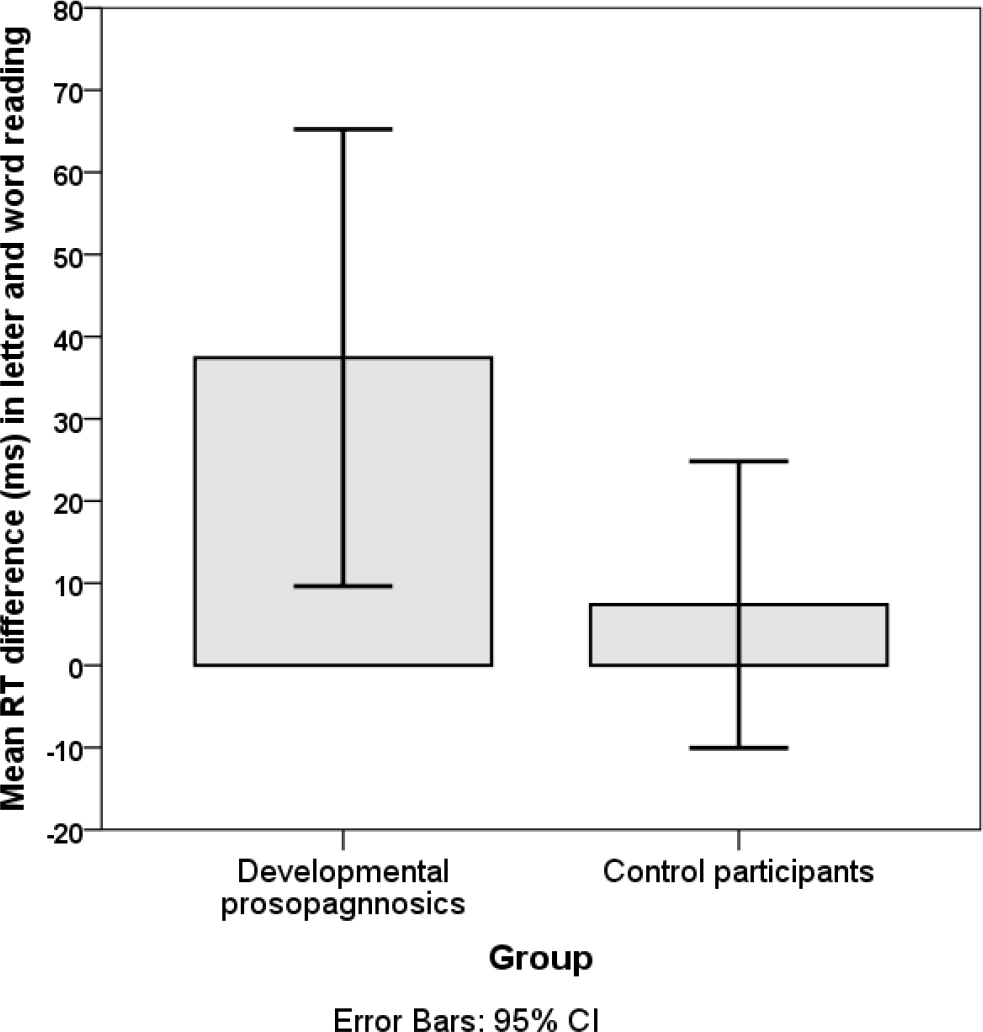
Word superiority in RTs. The mean difference in RT (ms) to letter and word naming for the DPs and control participants in Experiment 2a.

#### Experiment 2b. A psychophysical test of the word superiority effect

This experiment tested identification of briefly presented single stimuli (letters or words in different blocks) flashed at the centre of the screen, and followed by a pattern mask. There were 200 trials in each experimental block. In total, subjects ran 400 trials per stimulus type in an ABBA-design (letters first), and the first and second blocks for each stimulus type were preceded by 30 and 15 practice trials, respectively. In each trial, a single stimulus (word or letter) was chosen randomly and presented for one of ten exposure durations (10-100 ms, randomly intermixed). The stimulus was terminated by a pattern mask shown for 500 ms. Participants were instructed to make an unspeeded report of the stimulus, if they were “fairly certain” of its identity. Responses were recorded by the experimenter. To ensure foveal presentation, participants were required to focus on a centrally placed cross and then initiate the trial by pressing the left mouse button.

#### Results

We first compared the proportion of correct responses averaged across the ten exposure durations for the two groups for letters and words respectively.

The average proportion correct responses across all exposure durations was 0.74 (95% CI = [0.66, 0.82]) for letters, and 0.85 (95% CI = [0.82, 0.88]) for words in the DP-group. Hence, the WSE was large for the DPs (*d_z_* = 1.27; 95% CI *Mdif* = [0.04, 0.17]). For the controls, the average proportion correct responses across all exposure durations were 0.74 (95% CI = [0.69, 0.78]) for letters, and 0.82 (95% CI = [0.80, 0.85]) for words. Accordingly, the WSE was also large for the control participants (*d_z_* = 1.33; 95% CI *M_dif_* = 0.05, 0.12]). As can be seen from Figure 4 there is considerable overlap in the CIs for the DP-group and the control participants for both letters and words, suggesting that they also perform quite similarly in this experiment.

**Figure 4.**
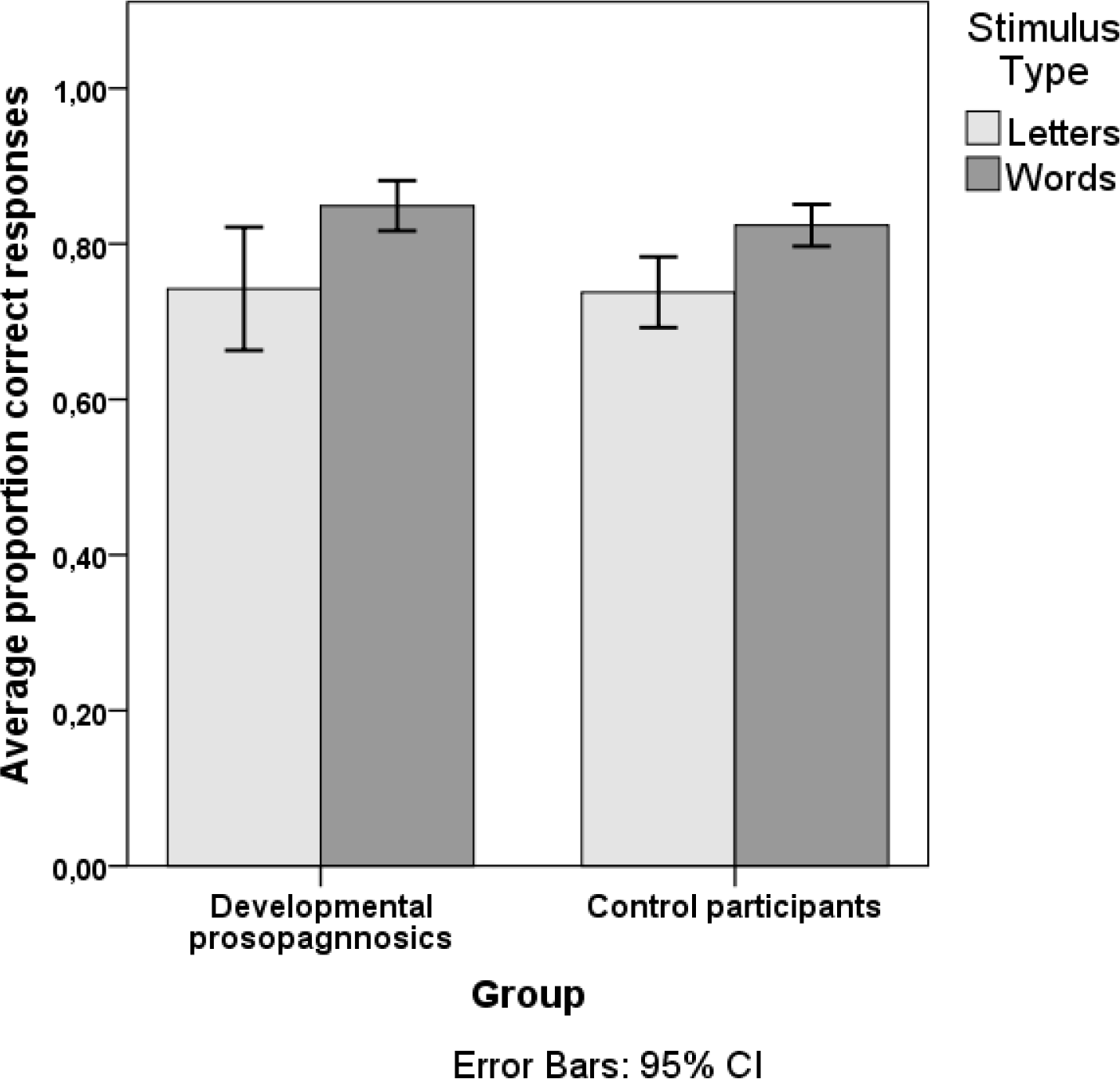
Experiment 2b. Shows the overall proportion correct responses for briefly presented words and single letters for DPs (*n* = 9) and matched controls (*n* = 18). The mean exposure duration for both groups was 55 ms.

We then compared the mean proportion of correct responses for letters and words for each of the ten exposure durations. Because four DPs received a slightly different set of exposure durations (caused by a computer problem resulting in a lower screen refresh rate), only five DPs (PP04, PP07, PP09, PP17 & PP27) and their matched controls *(n* = 10) were included in this analysis. As can be seen from Figure 5a (DPs) and Figure 5b (controls) the two groups performed quite similarly across the ten exposure durations. Performance generally increases with increased exposure duration, and the WSE is mainly evident at exposure durations between 20 and 60 ms for both groups. The variation (indexed by the CIs) appears somewhat larger for the DPs compared with the control participants at exposure durations shorter than 80 ms. This, however, could be a trivial consequence of the difference in sample size.

**Figure 5.**
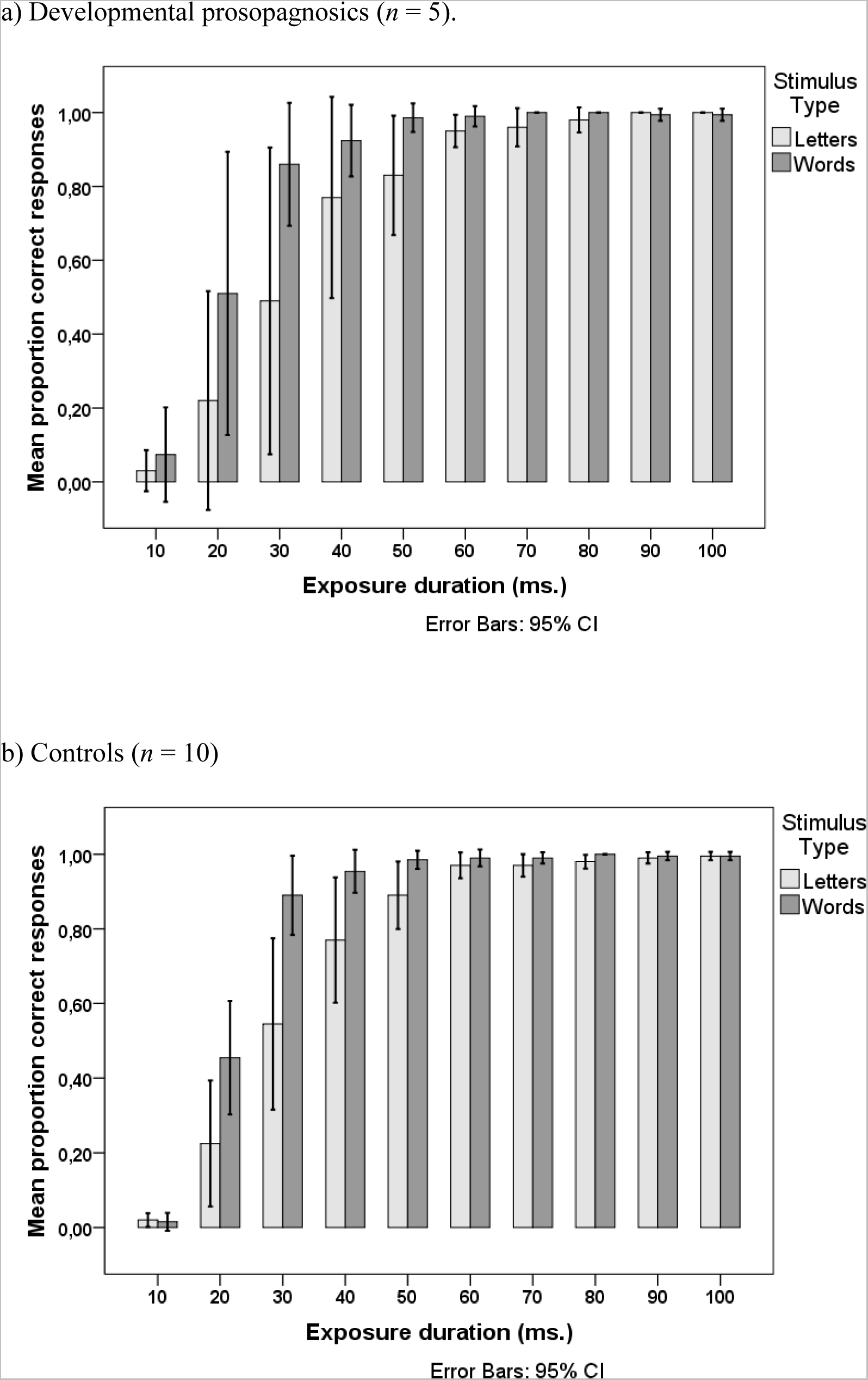
Mean proportion correct responses at the ten exposure durations for words and letters, in the DP-group (panel a) and controls (panel b) in Experiment 2b.

### Experiment 3. Text reading

#### Stimuli and procedure

The test was adapted from the standard 9 grade Danish reading tests (see Nielsen & Wilms, 2015 for a full description), and the main measure is reading time (in seconds). A text of 637 words was presented on the screen of a laptop computer. Instructions were to read the text carefully, as one would be required to answer questions about the text immediately after reading it. The text was a popular scientific text from a biology book^1^. This is the only one of our tests that had not been used in previous studies, and for that reason the procedure had some weaknesses (and was not entirely comparable to that described in Nielsen & Wilms, 2015). The text was too long to be shown on one page on a laptop, and part of the reading speed estimate is thus related to how long it took participants to move forward to the next page using the computer mouse. We bring the results here to show that despite this administration of the test, the results are very clear.

#### Results

Two of the DPs (PP04 and PP16) and two control participants (one for PP09 and one for PP10) did not perform the test. Hence, the DP group comprised 8 subjects and the control group 14 subjects.

In terms of accuracy on the text comprehension questions (four multiple choice questions), the DPs obtained a mean score of 2.5 (95% CI = [1.87, 3.13]) and the control participants a mean score of 2.64 (95% CI = [2.16, 3.13]). The mean reading speed of the DPs was 213 s (95% CI = [178, 247]) and 221 s (95% CI = [191, 251]) for the control participants. Hence, the performance of DPs and the control participants was quite alike in terms of both speed and comprehension.

## Discussion

Our findings are clear; our subjects with developmental prosopagnosia show no deficits in reading on the various tests included: Their letter and word naming times are similar to controls, they show a normal word superiority effect, a normal absence of a word length effect in single word reading, and normal text reading speed. As such, a clear dissociation between impaired face processing and preserved reading is evident in this group of developmental prosopagnosics.

Does this mean that reading and face recognition are completely independent processes, and that one function can be developmentally impaired without affecting the other? At least this seems true for one side of the dissociation; reading is intact in developmental prosopagnosia. A first glance at the literature also indicates that face recognition and naming may be intact in developmental dyslexia (Russeler, Johannes, & Munte, 2003; Smith-Spark & Moore, 2009), suggesting a double dissociation between reading and face processing. However, there are also reports of deficits in various face tasks, as well as different brain activation patterns for faces in dyslexics (Monzalvo, Fluss, Billard, Dehaene, & Dehaene-Lambertz, 2012; Tarkiainen, Helenius, & Salmelin, 2003). A very recent study has shown that adult dyslexics performed significantly below a matched control group on the Cambridge Face Memory Task (also used in our study), as well as other measures of face processing (Sigurdardottir, Ivarsson, Kristinsdottir, & Kristjansson, 2015).

This indicates that the relationship between reading and face processing is more complex than reading = left hemisphere, and faces = right hemisphere, and as reviewed in the introduction, there may be a developmental trajectory for hemispheric specialization that is at least partly caused by learning to read (Behrmann & Plaut, 2013; Cantlon et al., 2011; Dundas et al., 2014; Dundas, Plaut, & Behrmann, 2015). The “neuronal recycling” hypothesis (Dehaene & Cohen, 2011; Dehaene et al., 2010) is so far the most specific regarding the relationship between the two processes. This hypothesis suggests that when we learn to read, some brain areas originally contributing to face processing are “recycled” to be used in visual word recognition, and results supporting this come from studies of both children and illiterates learning to read (Cantlon et al., 2011; Dehaene et al., 2010). Our results show that normal visual word recognition can be acquired in spite of severe face recognition deficits, and at first glance this seems to contradict the “recycling” hypothesis. If it is correct that face recognition is bilaterally distributed from birth, but becomes more right lateralised with reading acquisition, then how might this work in DP to leave their reading functions unaffected? One possibility is that DP is a disorder characterised by cerebral dysfunction primarily in the posterior right hemisphere, perhaps parallel to the left hemisphere dysfunction seen in developmental dyslexia (Norton, Beach, & Gabrieli, 2015). This dysfunction in prosopagnosia thus affects the face processing network in the right hemisphere only or primarily, leaving the left hemisphere part of the network unaffected. The mechanism for recycling when learning to read, however, may be so strong, that regardless of the outcome for other functions (i.e. face processing), it may recycle the left hemisphere face processing areas for the purpose of visual word recognition. For prosopagnosics, then, learning to read recycles away what face recognition skills they may have had, to the benefit of reading skills. If this is the case, we should see a degradation in the face processing of DP’s in their early years of reading instruction. This is a clear hypothesis for future studies to test.

Another possibility is that the neural processes deficient in DP are localised more anteriorly in the face processing network, and do not implicate the fusiform areas affected by learning to read. Behrmann & Plaut (2013) review evidence supporting such a claim. In their “many-to-many” account, they propose that many posterior regions are necessarily engaged in the representation of multiple visual stimulus classes including words and faces, and that these regions form distributed but integrated large-scale circuits. If the fusiform regions involved in face processing are intact in DP, then the cerebral competition between reading and face processing in these regions might also proceed normally. This would result in the pattern of performance we report, impaired face recognition (due to cerebral dysfunction outside the posterior fusiform regions), and normal reading. The many-to-many account is partly based on findings that reading is affected in acquired prosopagnosia, and face recognition affected in pure alexia (Behrmann & Plaut, 2014), taken as evidence for shared, distributed networks being involved in both reading and face recognition. A recent study of acquired prosopagnosia has, however, suggested a modification of this claim: Hills, Pancaroglu, Duchaine, & Barton (2015) showed that patients with prosopagnosia due to unilateral right hemisphere damage did not have impaired word recognition, evidenced by normal RTs and WLEs. They did, however, show deficits in discriminating font or handwriting, which may suggest a hemispheric division of labour somewhat different from that originally suggested by Behrmann & Plaut (2013). Regardless of the neural processes involved, it is intriguing from a learning perspective that the developmental prosopagnosics can learn to read as fluently as normal subjects, while they are seemingly unable to learn efficient strategies for recognizing faces.

So what kind of process(es) is it that face recognition requires that word recognition does not? An answer to this question would take us considerably further in unravelling what might be the cause of developmental prosopagnosia. There is ample evidence suggesting that face recognition is particularly dependent on some kind of holistic or configural processing, which requires not only the identification of the parts (eyes, nose etc.), but also the specific relations between these parts; the configuration (Tanaka & Gordon, 2011). The same is true for efficient object recognition, especially when fine grained discriminations are required (Gerlach, 2009). In the case of object recognition we have argued that shape configuration – the binding of visual elements into elaborate shape descriptions in which relationships between the parts are specified – is augmented by global shape characteristics (the outline/gestalt of the object) (Gerlach, 2009). Importantly, these global shape characteristics are likely to be perceived *prior* to the object’s parts. A similar case has been made for face recognition where global shape characteristics also seem particularly important for identifying the specific configuration of the face parts (Goffaux, Hault, Michel, Vuong, & Rossion, 2005). For words, however, global shape does not seem to matter; we read words by recognizing the letters in parallel (parts first) and not by identifying the outline/gestalt (Grainger & Whitney, 2004; Pelli, Farell, & Moore, 2003). Hence, for faces and objects the configuration of parts and part identification usually follows after perception of the whole object. For words, recognition is a product of part (letter) perception. Based on these considerations it seems plausible that the face recognition problems in our sample of DPs could be caused (at least partly) by impaired processing of global shape. As global shape plays no part in word recognition, reading is intact in these individuals despite the fact that word recognition is a perceptually demanding task.

So reading and face recognition are indeed different, and face recognition can be impaired while reading, and the ability to learn to read, is unaffected. Perhaps Oliver Sacks could have told us so from the beginning. While he never described his reading abilities in as much detail as his problems with face recognition, at least it is evident that he wrote better than most…

## Footnotes

1 One DP-participant was presented with a different text from the same book, matched in length and difficulty, by mistake.

## Acknowledgements

We wish to thank the participating subjects, Fakutsi for face value, and Simon Nielsen for providing the text reading test.

## Funding acknowledgment

This project was supported by a grant from the Danish Research Council for Independent Research | Humanities (grant no. DFF – 4001-00115) to C.G. and R.S.

**Supplementary Table S1.**
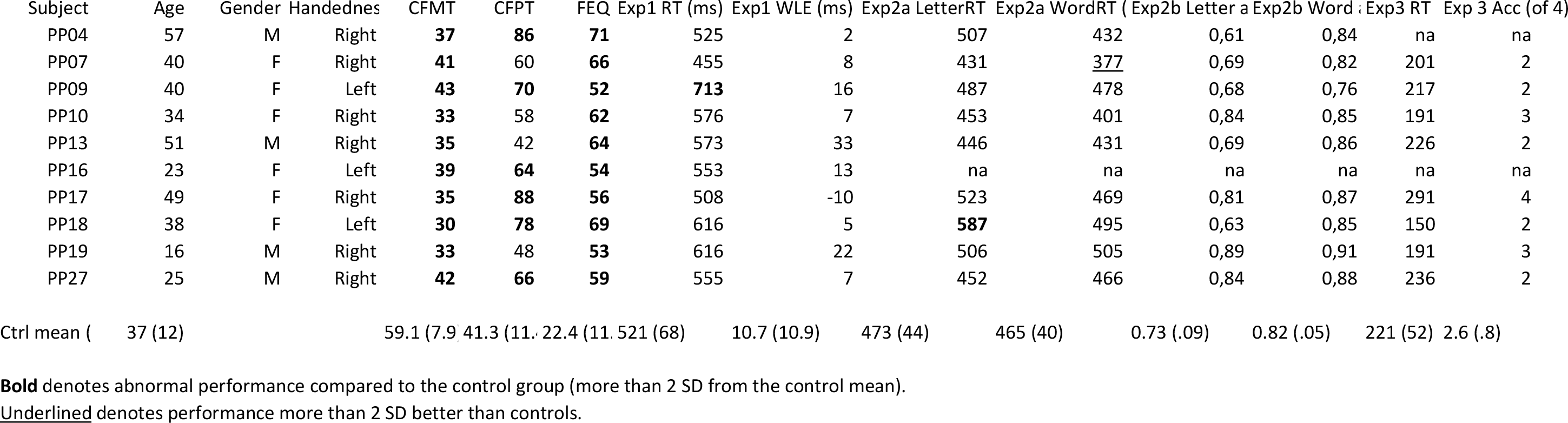
Individual experimental data.

## References

1. Baron-Cohen, S., Wheelwright, S., Skinner, R., Martin, J., & Clubley, E. (2001). The autism-spectrum quotient (AQ): evidence from Asperger syndrome/high-functioning autism, males and females, scientists and mathematicians. Journal of autism and developmental disorders, 31 (1), 5–17.

2. Barton, J. J., S., Hanif, H. M., Björnström, L. E., & Hills, C. (2014). The word length effect in reading: A review. Cognitive Neuropsychology (31 (5-6)), 378–412.

3. Behrmann, M., & Plaut, D. C. (2013). Distributed circuits, not circumscribed centers, mediate visual recognition. Trends in cognitive sciences, 17 (5), 210–219. doi: http://dx.doi.org/10.1016/j.tics.2013.03.007

4. Behrmann, M., & Plaut, D. C. (2014). Bilateral Hemispheric Processing of Words and Faces: Evidence from Word Impairments in Prosopagnosia and Face Impairments in Pure Alexia. Cerebral Cortex, 24 doi: doi: 10.1093/cercor/bhs390

5. Bowles, D. C., McKone, E., Dawel, A., Duchaine, B., Palermo, R., Schmalzl, L.,… Yovel, G. (2009). Diagnosing prosopagnosia: effects of ageing, sex, and participant-stimulus ethnic match on the Cambridge Face Memory Test and Cambridge Face Perception Test. Cognitive Neuropsychology, 26 (5), 423–455. doi: 10.1080/02643290903343149

6. Cantlon, Jessica F., Pinel, Philippe, Dehaene, Stanislas, & Pelphrey, Kevin A. (2011). Cortical Representations of Symbols, Objects, and Faces Are Pruned Back during Early Childhood. Cerebral Cortex (New York, NY), 21 (1), 191–199. doi: 10.1093/cercor/bhq078

7. Cohen, L., Dehaene, S., Naccache, L., Lehericy, S., Dehaene-Lambertz, G., Henaff, M. A., & Michel, F. (2000). The visual word form area: spatial and temporal characterization of an initial stage of reading in normal subjects and posterior split-brain patients. Brain, 123 (Pt 2), 291–307.

8. Cumming, G. (2014). The new statistics: why and how. Psychol Sci, 25 (1), 7–29. doi: 10.1177/0956797613504966

9. Dehaene, S., & Cohen, L. (2007). Cultural recycling of cortical maps. Neuron, 56 (2), 384–398. doi: 10.1016/j.neuron.2007.10.004

10. Dehaene, S., & Cohen, L. (2011). The unique role of the visual word form area in reading. Trends in Cognitive Science, 15 (6), 254–262. doi: 10.1016/j.tics.2011.04.003

11. Dehaene, Stanislas, Pegado, Felipe, Braga, Lucia W, Ventura, Paulo, Nunes Filho Gilberto, Jobert, Antoinette, … Cohen, Laurent. (2010). How learning to read changes the cortical networks for vision and language. Science, 330 (6009), 1359–1364.

12. Duchaine, B., Germine, L., & Nakayama, K. (2007). Family resemblance: ten family members with prosopagnosia and within-class object agnosia. Cognitive Neuropsychology, 24 (4), 419–430. doi: 10.1080/02643290701380491

13. Duchaine, B., & Nakayama, K. (2006). The Cambridge Face Memory Test: results for neurologically intact individuals and an investigation of its validity using inverted face stimuli and prosopagnosic participants. Neuropsychologia, 44 (4), 576–585. doi: 10.1016/j.neuropsychologia.2005.07.001

14. Dundas, E. M., Plaut, D. C., & Behrmann, M. (2013). The joint development of hemispheric lateralization for words and faces. Journal of Experimental Psychology General, 142 (2), 348–358. doi: 10.1037/a0029503

15. Dundas, E. M., Plaut, D. C., & Behrmann, M. (2014). An ERP investigation of the co-development of hemispheric lateralization of face and word recognition. Neuropsychologia, 61, 315–323. doi: 10.1016/j.neuropsychologia.2014.05.006

16. Dundas, E. M., Plaut, D. C., & Behrmann, M. (2015). Variable Left-hemisphere Language and Orthographic Lateralization Reduces Right-hemisphere Face Lateralization. Journal of cognitive neuroscience, 27 (5), 913–925. doi: 10.1162/jocn_a_00757

17. Gazzaniga, M. S., Ivry, R. B., & Mangun, G. R. (2013). Cognitive neuroscience: the biology of the mind (Fourth edition. ed.). New York, N.Y.: W. W. Norton & Company, Inc.

18. Gerlach, C. (2009). Category-specificity in visual object recognition. Cognition, 111 (3), 281–301. doi: 10.1016/j.cognition.2009.02.005

19. Goffaux, V., Hault, B., Michel, C., Vuong, Q. C., & Rossion, B. (2005). The respective role of low and high spatial frequencies in supporting configural and featural processing of faces. Perception, 34 (1), 77–86.

20. Grainger, J., & Whitney, C. (2004). Does the huamn mnid raed wrods as a wlohe? Trends in Cognitive Science, 8 (2), 58–59. doi: 10.1016/j.tics.2003.11.006

21. Habekost, T., Petersen, A., Behrmann, M., & Starrfelt, R. (2014). From word superiority to word inferiority: visual processing of letters and words in pure alexia. Cognitive Neuropsychology, 31 (5-6), 413–436. doi: 10.1080/02643294.2014.906398

22. Hills, Charlotte S., Pancaroglu, Raika, Duchaine, Brad, & Barton, Jason J. S. (2015). Word and text processing in acquired prosopagnosia. Annals of Neurology, 78 (2), 258–271. doi: 10.1002/ana.24437

23. Kanwisher, N., McDermott, J., & Chun, M. M. (1997). The fusiform face area: a module in human extrastriate cortex specialized for face perception. Journal of Neuroscience, 17 (11), 4302–4311.

24. Kennerknecht, I., Grueter, T., Welling, B., Wentzek, S., Horst, J., Edwards, S., & Grueter, M. (2006). First report of prevalence of non-syndromic hereditary prosopagnosia (HPA). American Journal of Medical Genetics Part A, 140 (15), 1617–1622. doi: 10.1002/ajmg.a.31343

25. Monzalvo, K., Fluss, J., Billard, C., Dehaene, S., & Dehaene-Lambertz, G. (2012). Cortical networks for vision and language in dyslexic and normal children of variable socio-economic status. NeuroImage, 61 (1), 258–274. doi: 10.1016/j.neuroimage.2012.02.035

26. Nielsen, Simon, & Wilms, Inge Linda. (2015). Cognitive ageing on latent constructs for visual processing capacity: A novel Structural Equation Modelling framework with causal assumptions based on A Theory of Visual Attention. Frontiers in Psychology, 5. doi: 10.3389/fpsyg.2014.01596

27. Norton, E. S., Beach, S. D., & Gabrieli, J. D. (2015). Neurobiology of dyslexia. Current opinion in neurobiology, 30, 73–78. doi: 10.1016/j.conb.2014.09.007

28. Pelli, D. G., Farell, B., & Moore, D. C. (2003). The remarkable inefficiency of word recognition. Nature, 423 (6941), 752–756. doi: 10.1038/nature01516

29. Reicher, G. M. (1969). Perceptual recognition as a function of meaninfulness of stimulus material. Journal of experimental psychology, 81 (2), 275–280.

30. Russeler, J., Johannes, S., & Munte, T. F. (2003). Recognition memory for unfamiliar faces does not differ for adult normal and dyslexic readers: an event-related brain potential study. Clin Neurophysiol, 114 (7), 1285–1291.

31. Sacks, O. (2010). Face-blind: why are some of us terrible at recognizing faces? New Yorker, 36–43.

32. Sacks, O. W. (1985). The man who mistook his wife for a hat and other clinical tales. New York: Summit Books.

33. Sigurdardottir, H. M., Ivarsson, E., Kristinsdottir, K., & Kristjansson, A. (2015). Impaired Recognition of Faces and Objects in Dyslexia: Evidence for Ventral Stream Dysfunction? Neuropsychology. doi: 10.1037/neu0000188

34. Smith-Spark, J. H., & Moore, V. (2009). The representation and processing of familiar faces in dyslexia: differences in age of acquisition effects. Dyslexia, 15 (2), 129–146. doi: 10.1002/dys.365

35. Starrfelt, R., Nielsen, S., Habekost, T., & Andersen, T. S. (2013). How low can you go: Spatial frequency sensitivity in a patient with pure alexia. Brain and Language, 126 (2), 188–192. doi: http://dx.doi.org/10.1016/j.bandl.2013.05.006

36. Starrfelt, R., Petersen, A., & Vangkilde, S. (2013). Don’t words come easy? A psychophysical exploration of word superiority. Frontiers in Human Neuroscience, 7, 519. doi: 10.3389/fnhum.2013.00519

37. Tanaka, J. W., & Gordon, I. (2011). Features, Configuration, and Holistic Face Processing. In A. J. Calder, G. Rhodes, M. H. Johnson & J. V. Haxby (Eds.), The Oxford Handbook of Face Perception (pp. 177 – 194). New York: Oxford University Press.

38. Tarkiainen, A., Helenius, P., & Salmelin, R. (2003). Category-specific occipitotemporal activation during face perception in dyslexic individuals: an MEG study. Neuroimage, 19 (3), 1194–1204.

39. Wheeler, D. D. (1970). Processes in word recognition. Cognitive Psychology, 1 (1), 59–85. doi: http://dx.doi.org/10.1016/0010-0285(70)90005-8

